# Scaling laws in spoken language associated with cognitive functions

**DOI:** 10.1101/841247

**Authors:** Masato S. Abe, Mihoko Otake-Matsuura

## Abstract

A decline in cognitive functions due to aging has led to critical problems in modern society, and it is imperative to develop a method to predict the decline or related diseases, including dementia. Although it has been expected that language could be a sign of the cognitive decline, it remains less understood, especially in natural conditions. In this study, we recorded the large-scale data of one million words from group conversations among healthy elderly people and analysed the relationship between spoken language and cognitive functions based on scaling laws, namely, Zipf’s law and Heaps’ law. We found that word patterns followed these scaling laws irrespective of cognitive function, but the variations in scaling laws were associated with cognitive functions. Moreover, using generative models, we uncovered the relationship between the variations and cognitive functions. These results indicate that scaling laws in language can be a biomarker for the cognitive decline.

## Main

Understanding the aging of brain functions and predicting cognitive decline are crucial for modern aging societies because mental health problems in elderly people have a huge impact on their daily life and are causing significant medical and economic costs in many countries around the world^1,2^. The most common example of cognitive decline with age is dementia, especially Alzheimer’s disease, which leads to the impairment of the executive functions or behavioural and psychological symptoms^3,4^. While the neuropathological mechanisms for dementia are considered as remarkably complex^2^, it is necessary to urgently develop practical methods to predict the decline or maintain healthy cognitive functions. It is expected that language may be a key sign representing states of cognitive functions^5–9^.

Language is the most sophisticated means of communication for humans; it allows abstract thoughts and underlies our various social activities ranging from daily conversations to cultural accumulation^10,11^. The complex language processing clearly comes from our brain functions; thus, the impairments in cognitive functions can result in language disorder^12,13^. Therefore, it is imperative to understand the relationship between language and aging in predicting cognitive functions using data related to language. Some methods for predicting cognitive diseases using language data have been proposed^5–9,12–14^. For instance, scores of verbal fluency tests (in which participants produce as many words as possible from a category in 60 seconds) are one of the indices used to distinguish people with dementia or mild cognitive impairment (MCI) from healthy people^12^. Recently, it has become easier to record and analyse large-scale data on language such as ordinary conversations due to the development of devices and algorithms^7–9^. Therefore, we expect that extracting information about cognitive functions from large-scale data makes it possible to develop a more prominent method. However, the understanding of the relationship between cognitive functions and language in natural conditions such as ordinary conversations is still lacking. In this study, we focused on statistical laws, namely, the scaling laws of spoken language in conversations.

Scaling laws, which are generally defined as *f*(*x*) ∼ *x*^*µ*^, where *µ* is a scaling exponent (here, “∼” means that the left side is proportional to the right side), are ubiquitous in natural phenomena^15^. Interestingly, they are observed widely in phenomena related to brains or behaviour, such as neural dynamics, decision-making, semantic memory, memory retrieval, cognition, movements, language, and social dynamics^16–24^. The most famous example can be seen in the word patterns of human language^16^. So far, the previous studies that have focused on language patterns in corpus data from written texts or spoken language^16,19,25,26^ have found two main scaling laws, namely, Zipf’s law and Heaps’ law. Zipf’s law states that the frequency of appearance of words with rank *r* for appearance follows a kind of power-law distributions *P*(*r*) ∼ *r*^−*α*^, suggesting that a huge number of words are rarely used, while a small number of words are extremely used often. Since it was reported that the exponent *α* is close to 1 in most cases^27^, the most frequent word will appear twice as often as the second most frequent word, three times as often as the third one, and so on.

Heaps’ law, denoted as *N* ∼ *M*^*β*^, indicates that the number of different words *N* (i.e. types) sub-linearly increases as the number of words *M* (i.e. tokens) increases^28^. In other words, Heaps’ law describes how new words are produced along with sentences or during conversations. Empirical studies on written or spoken language have reported that the exponent *β* is estimated to be about 0.7^25,26^. Note that it has been indicated that Zipf’s and Heaps’ laws were universal, regardless of language, even though the human language has highly complex structures in terms of context or grammar^25–27^.

Most studies on these scaling laws have been conducted from statistical and theoretical standpoints^16,25,29,30^. It is unclear whether the spoken language in elderly people with low cognitive functions follow these scaling relationships or not and how the variations in these scaling relationships are related to cognitive function, although the Zipf’s laws in children and adults and in schizophrenia cases were reported^31–33^. Moreover, some proposed models can produce the scaling laws, but most of them lack realistic mechanisms such as the psychological process, memory, and so on^27^. Hence, it is still not fully understood how and why these scaling laws emerge in natural language.

In the present study, we focused on the spoken language of healthy elderly people with various cognitive functions under natural conditions. The questions included i) whether natural spoken language in elderly people follows the scaling laws mentioned above or not and, ii) if so, how the variations in scaling laws are related to cognitive scores. The study revealed the relationship between the scaling laws and cognitive functions. Based on the findings, we propose the scaling exponents as a biomarker for detections of early cognitive decline.

## Results

### Zipf’s law and Heaps’ law in spoken language

We recorded conversations among healthy elderly participants (*n* = 65) and conducted independent cognitive tests to evaluate cognitive functions in advance of overall conversational experiments (the details in Methods). First, we show the basic characteristics of conversations among participants in Fig S1. The mean of total spoken words among the participants was 18,066 ± 7,213 (SD), and the mean of different words was 2,308 ± 519 (SD). Fig. 1a shows the rank-frequency distribution of words and Fig. 1b shows the relationship between the number of words and the number of different words spoken by each participant, colouring the lines according to cognitive function scores. The rank-frequency distributions apparently follow a kind of power-law distribution because the frequencies of words, except for higher rank (*r* < 10∼), seem to be on the line in the log-log plot, suggesting that the spoken words of elderly people follow Zipf’s law. To verify this statistically, we fitted six candidate distributions, including a power-law distribution, a shifted power-law distribution, and so on to our empirical data using rigorous statistical techniques^25,34^ (see Methods). The results obtained by the model selection showed that all the rank-frequency distributions for the 65 participants were fitted to shifted power-law distributions (Akaike weights of shifted power-law distribution > 0.99). This statistical result suggests that spoken words in conversations among healthy elderly people follow Zipf’s law, although the words in the higher rank (i.e. small *r*) do not follow a pure power-law distribution. This is not surprising because it is consistent with previous results that written language followed a shifted power-law distribution^16,27^. The estimated Zipf’s exponents *α* were 1.22 ± 0.048 (mean ± SD), and the range was [1.11, 1.32].

**Fig. 1:**
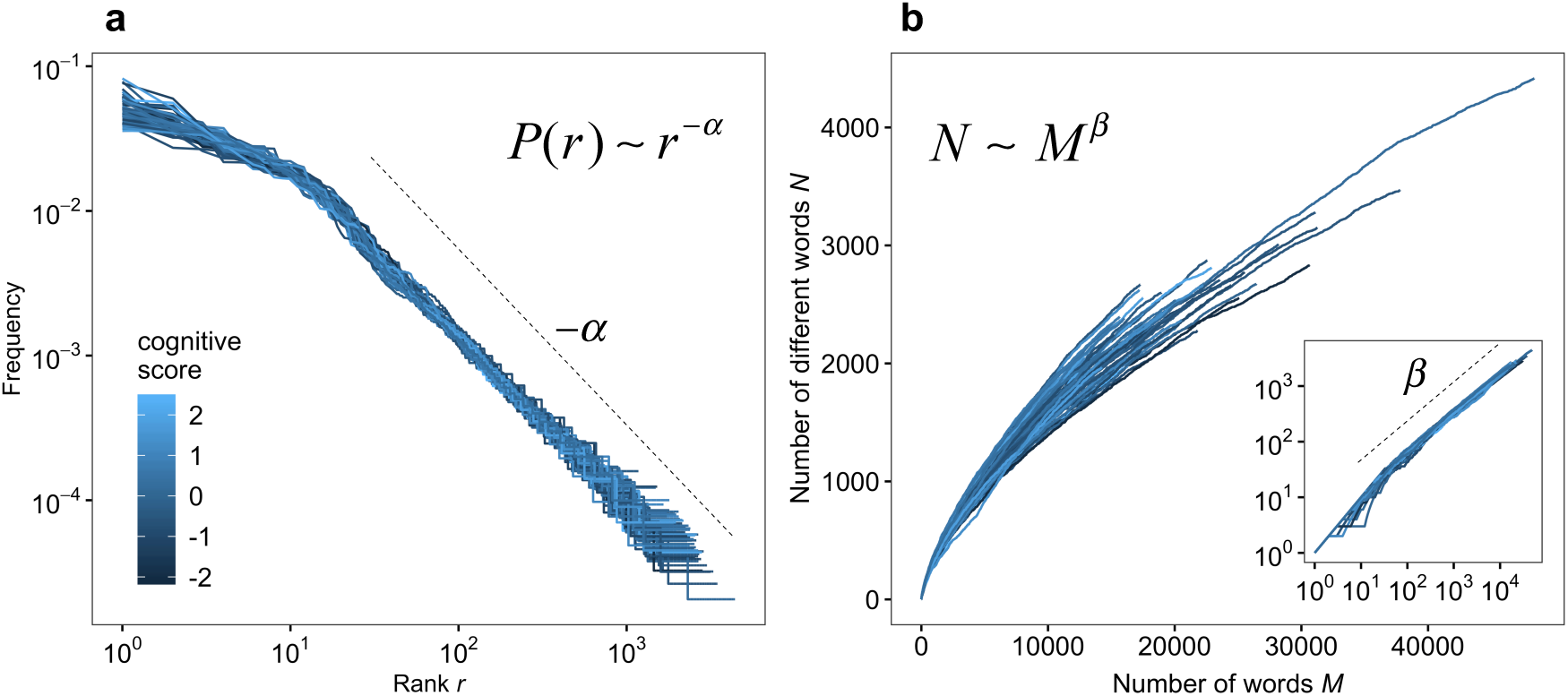
Zipf’s law and Heaps’ law in natural conversations of elderly people (a) Zipf’s law: relationship between rank *r* and frequency of words. The slope corresponds to the scaling exponent −*α*. (b) Heaps’ law: relationship between the number of words *M* and the number of different words *N*. The inset in (b) is a log-log plot of the main plot in (b) and, the slope corresponds to the scaling exponent *β*. In (a) and (b), each line represents each participant, and the colour represents their cognitive scores shown in (a).

Then, we analysed the relationship between the number of words and the number of different words. Based on the results, it seems to follow Heaps’ law because there is a straight line in the log-log plot (see inset in Fig. 1b). Furthermore, there seems to be a relationship with slope = 1 between *N* and *M* at small *M* (i.e. *M* < *m*, where *m* is a break point) because most words are new words in the vicinity of the initial states. Therefore, we fitted a double power-law model *N* = *M*^1^ (*M* < *m*) and *N* ∼ *M*^*β*^ (*M* ≥ *m*) to the data^25^, estimated both the scaling exponent *β* and threshold *m* using an optimisation method (Nelder-Mead method), and then calculated R-squared values to evaluate the good-of-fit of the model to the data. We found that the Heaps’ exponents *β* was 0.685 ± 0.02 (mean ± SD), the range was [0.637, 0.739], and the averaged R-squared value was 0.999 ± 0.001 (SD) for an overall range of *M*. This statistical analysis strongly reveals that the spoken words of elderly people also follow Heaps’ law. Moreover, the mean of the estimated break points was *m* = 32.

### Relationship between word production patterns and cognitive score

Next, we explored the relationships between the word production patterns including the scaling laws estimated above and the cognitive scores obtained from a cognitive test conducted independently. First, although one may expect that talkative people have high cognitive scores, we found no evidence of a relationship between the cognitive score and the total number of words spoken (Spearman’s correlation coefficient *ρ* = −0.11, *p* = 0.38). Second, we straightforwardly calculated a type-token ratio, which is defined as a value of the number of different words divided by the number of words. The Pearson’s correlation coefficient between the ratio and the cognitive score was 0.18 (*p* = 0.15), thus there was no significant relationship between the type-token ratio and the cognitive scores. Then, we analysed the relationship between the cognitive score and Zipf’s law and found no significant relationship (correlation coefficient *r* = −0.23, *p* = 0.07; Fig. 2a; Supplementary Table 3).

**Fig. 2:**
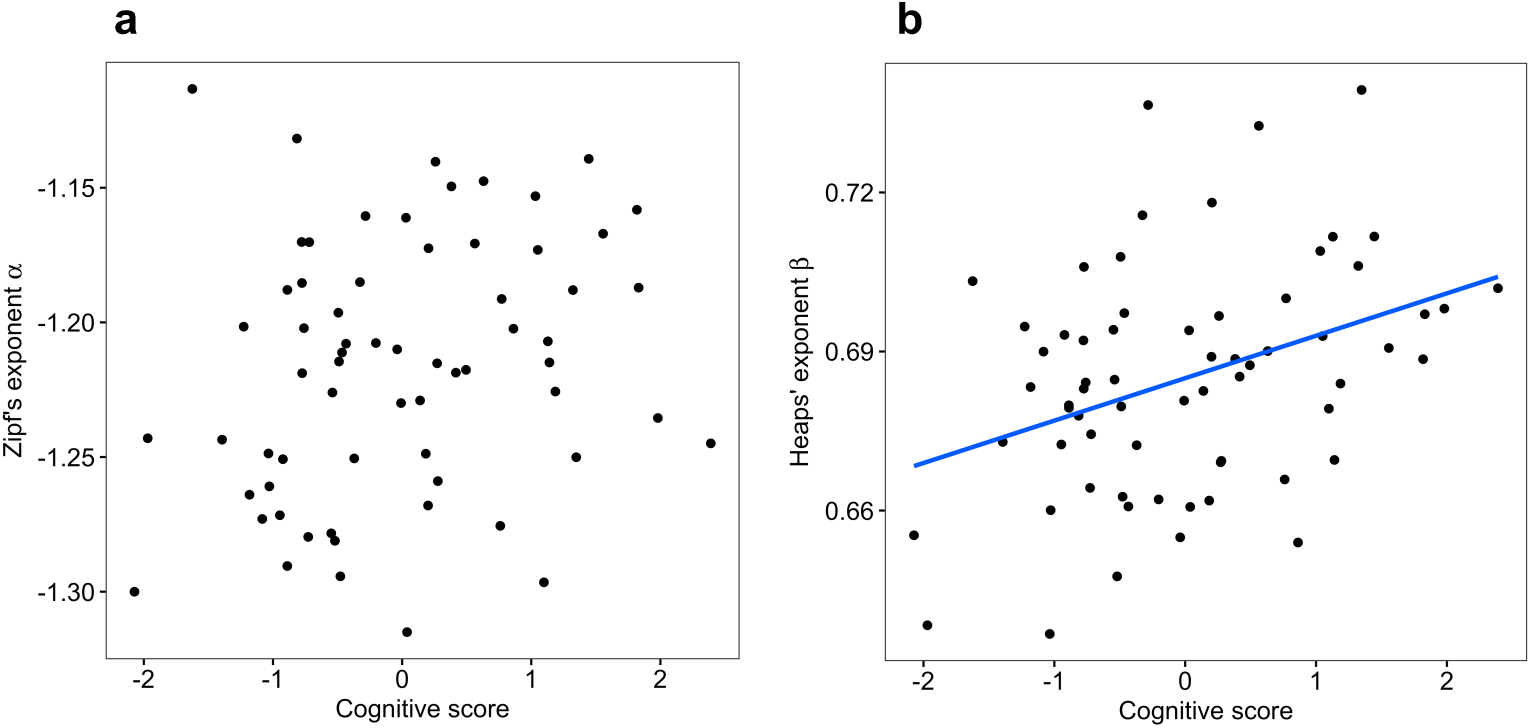
Relationship between scaling laws and cognitive score (a) Cognitive scores vs Zipf’s exponent *α*. Each point represents the data for each participant. (b) Cognitive scores vs Heaps’ exponent *β*. The blue line is the model derived from the best-fitted model, and the grey shade represents the 95% confidence interval.

Then, we examined the relationship between the exponent *β* of Heaps’ law and cognitive scores and we found a significant positive correlation between the exponents *β* and cognitive scores (Pearson’s correlation coefficient *r* = 0.37, *p* = 0.002; Fig. 2b). The result of the generalised linear model indicates that the coefficients of exponent *β* and gender were significant (*p* = 0.002 and *p* = 0.033, respectively; Supplementary Table 3). The participants with higher cognitive scores were likely to have word patterns with higher Heaps’ exponent *β*, and vice versa. On the other hand, the result for the type of conversation and the age of participants showed that they were not associated with the exponent (*p* = 0.75 and *p* = 0.72, respectively; Supplementary Table 3). Then, we confirmed that the relationship was robust to original cognitive scores. The correlation coefficients between the exponents and each of the four original cognitive scores (MoCA-J, logical memory I + II, digit symbol coding, digit span) were also significantly large (Supplementary Table 4). Thus, these results indicate that the variation of the Heaps’ law could come from the difference of the cognitive functions.

When one uses conversational data as a biomarker of cognitive decline, it is useful to investigate the relationship between the number of words (i.e. data length) and the degree of association of scaling laws with cognitive scores. We calculated the correlation coefficient between the exponents and cognitive scores with different data lengths. Fig. 3a shows that in Zipf’s law, no correlation was found, which is consistent with the result in the case that all data were used. In contrast, Fig. 3b shows that the longer the data length, the higher the correlation coefficient between Heaps’ exponent and cognitive score. Importantly, this indicates that it is not necessary to analyse data sets with tens of thousands of words of each participant. Therefore, the result suggests that we could quantify the association with cognitive scores with as little as utterances in an hour or two per person, suggesting that we can extract useful information about cognitive functions in realistic conditions such as ordinary conversations.

**Fig. 3:**
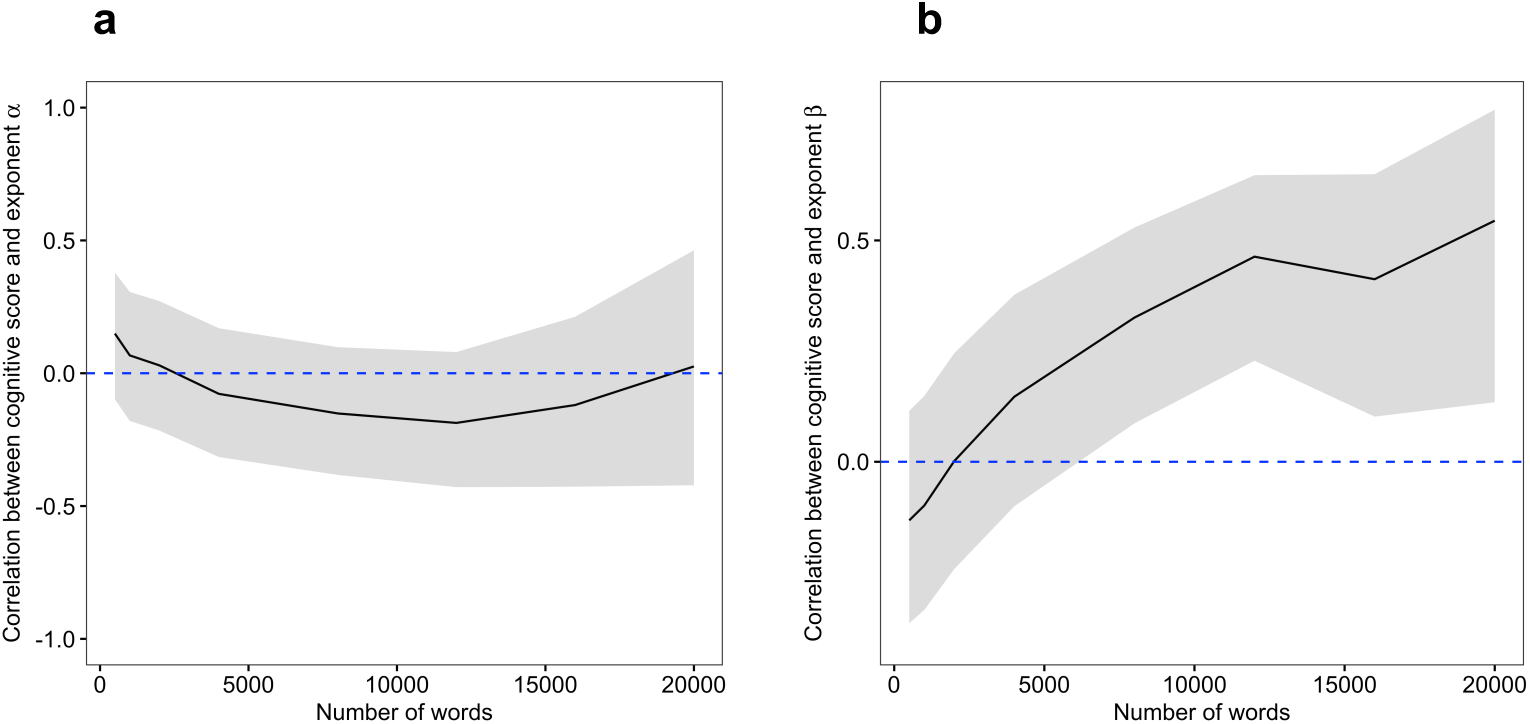
Relationship between data length (number of words) and correlation coefficients with exponent (a) and (b) correspond to Zipf’s law and Heaps’ law, respectively. The solid line represents Pearson’s correlation coefficient between cognitive scores and the exponents *α* or *β* when increasing the number of words, the shade is the 95% confidence interval, and the blue dashed line is 0.

### Generative models bridging scaling laws and cognitive functions

In this section, we focus on the Heaps’ exponent because we obtained the evidence of a significant association with cognitive functions. Next, we asked a question why the cognitive functions are associated with the Heaps’ exponent. To mechanically bridge them, we used a generative model for scaling laws in language developed by Gerlach and Altmann^25^, although various models have been proposed^29^. The model we used is a stochastic model originally based on the Yule process^35^, and can produce Zipf’s law and Heaps’ law from a simple assumption (see Methods). Here, we focused on a parameter related to the decay rate of probability for new word production and interpret it as a cognitive function. We analysed the relationship between the scaling exponent *β* and the parameters (see Method). We set the maximum number of words *M*_max_ = 20,000, which is close to the empirical data length and a relatively small value compared with the books corpus data (e.g. 10^9^ words in ^25^). Moreover, we fitted the double power-law model to simulated data and estimated the exponent. Fig. 4a shows the relationship between the number of words and different words derived from the model. When the parameter *γ* of cognitive functions changes, the patterns of Heaps’ law can change (Fig. 4a). Further, the parameters significantly correspond to the exponent *β*, and the one with the lower cognitive function can produce Heaps’ law with lower *β* (Fig. 4b), suggesting that the exponent *β* means that the growth rate of new words and the higher values correspond with high growth rate of new words, and vice versa. Note that the model and the relationship between the parameters and the exponent we observed are not novel, but we confirmed that the statistical fitting for simulated data from the model can recover the relationship even for the small number of words.

**Fig. 4:**
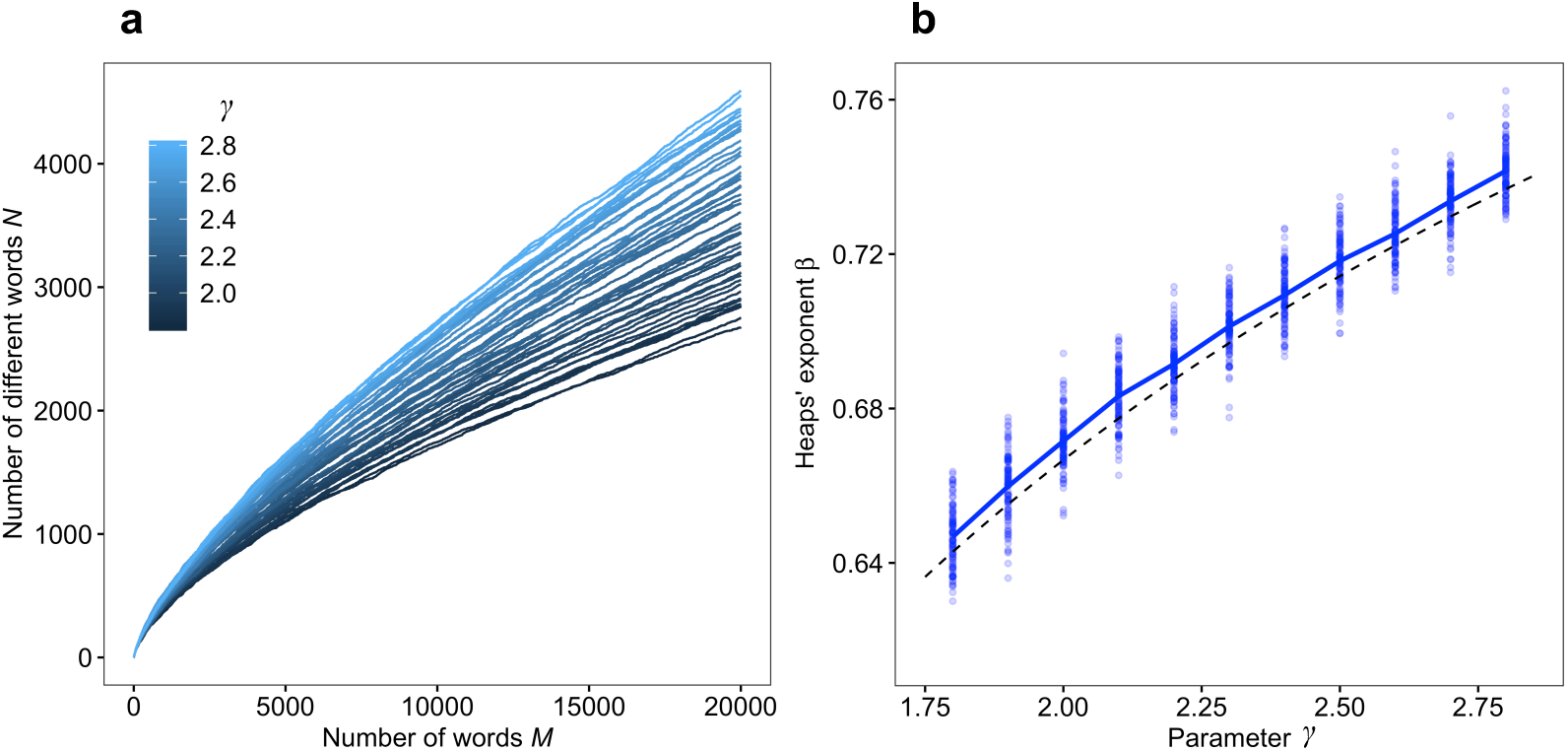
Computational model for words production (a) The result for simulation of word production. Each line represents one run of the simulation, and the colour represents the parameter *γ*. The other parameter *s* is set to 32, which was empirical averaged values of the break point of double power-law fitting. (b) The relationship between the parameter *γ* and the scaling exponent *β*. The blue solid line and points represent the mean and each data point of simulations, and the dashed line means the relationship obtained from the analytical solution.

### Source of new words

The mathematical model indicates that the scaling exponent of Heaps’ law could come from the production rate of new words. Therefore, knowledge about the source of the new words could provide new insights into how people produce new words. To investigate this, we extracted the information on where new words come from, namely, from the participants’ own internal memory or from other participants’ utterances during conversations. When a participant produced a novel word, we checked whether the words had already been used by the other group members by the time of the session. If the word had already been used, it was roughly considered that the participant had heard the word through conversation, which suggests that the participant took the new information from others. If not, there is a high possibility that the word came from the participant’s internal memory. To quantify the source of new words, we calculated the ratio of new words from other participants based on the total number of new words. Note that this analysis was conducted only for the free conversation group because participants in the presentation group were determined artificially and the order of speaking was heterogeneous among them. Fig. 5 shows that the high exponents of Heaps’ law are related to the high ratio of new words from other participants (*r* = 0.43, *p* = 0.01), suggesting that the high susceptibility to other participants can provide a new word production rate.

**Fig. 5:**
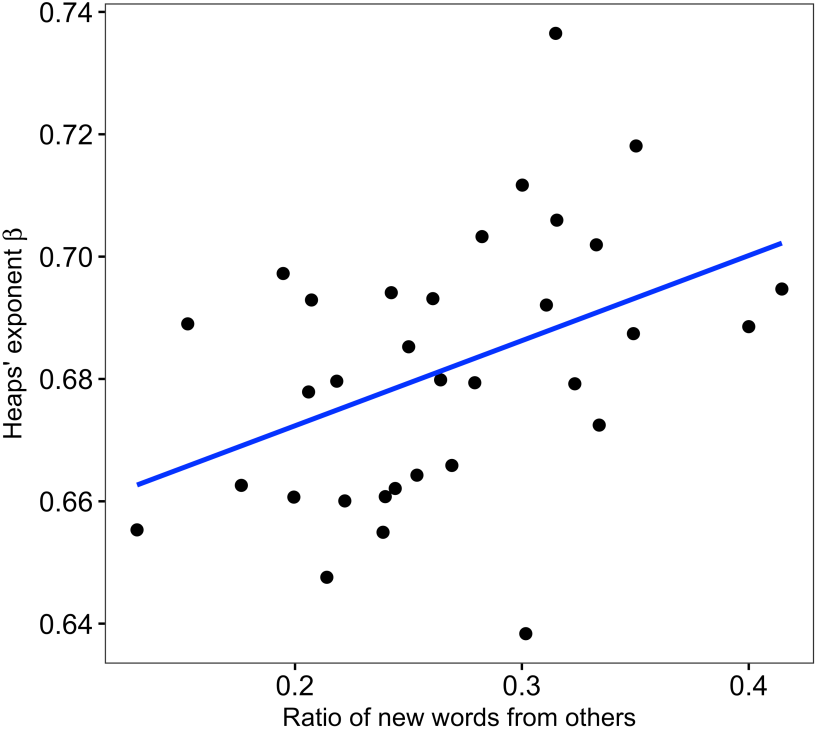
Source of new words and Heaps’ exponent *β* The new words can be classified into the words from one’s own memory and the words from others. The horizontal axis represents the ratio of the new words from others. Each point represents each participant in free conversation conditions. The blue line represents the best-fitted linear model of the relationship, and the shade is the 95% confidence interval.

## Discussions

In this study, we quantitatively investigated the natural spoken language of healthy elderly people with various cognitive function scores from the viewpoint of scaling laws in word patterns and explored the relationship between the scaling laws including Zipf’s law and Heaps’ law and cognitive function scores (Fig. 1). We found that the scaling laws in spoken language were robust, irrespective of various cognitive function scores from the result of fitting Zipf’s law and Heaps’ law. We did not find a significant relationship between the Zipf’s exponent and the cognitive score (Figs. 2a, 3a). The exponents of Heaps’ law, that is, the slope of relationship between the number of words and different words, were significantly associated with cognitive function scores (Figs. 2b, 3b). The relationship with Heaps’ law was supported by the stochastic model for word production patterns (Fig. 4). Moreover, the large Heaps’ exponents were related to obtaining new words from others and using them in their conversations (Fig. 5). Note that the participants were healthy elderly people after screening using criteria MMSE score ≥ 24, although there were variations in their cognitive scores. Therefore, we can exploit the relationship with scaling laws as a biomarker to detect the tendency of cognitive decline, even for people regarded as being healthy.

A feature of our approach for conversational data is that large-scale data make it possible to analyse statistical patterns such as scaling laws, which have been considered as robust properties across language and cultures^36^. Therefore, although the participants in our study were Japanese, our results and implications could be extended to other languages. Furthermore, by focusing only on the statistical patterns and scaling laws, we can exclude the meaning of words or contents of a conversation and separate the frame (i.e. scaling function) and the variations (i.e. exponent). This is considered as a type of a coarse-graining method for large and complex data while keeping important information, which could extract understandable characteristics from large data of language. In this sense, our approach is different from machine learning approaches that specialise in prediction rather than understanding the mechanisms behind the word patterns. Thus, it is considered that our approach does not need much data and can detect the relationship between cognitive functions and the word patterns even for healthy participants.

Our findings suggest that healthy elderly people with variations of cognitive scores are still on the scaling law (Fig. 1). In contrast, MCI or dementia patients might not be on the scaling laws because repeating a certain word due to critical cognitive impairment or memory disorder may result in the collapse of the scaling laws. Previous studies have reported that the Zipf’s exponents of schizophrenia patients are different from those of healthy people^31^. However, the rigorous statistical techniques that we used in the present study for fitting power-law distribution were developed recently; thus, they can reveal the patterns of scaling laws more accurately^34^. Hence, future studies should include analysing whether the word patterns in patients with cognitive disorders or other mental disorders follow scaling laws or not, and how different their exponents are. This can help in understanding why and how the scaling laws emerge and the development of early detection methods for the disorders.

The production of new words is important for communication or creating new ideas. There would be two points regarding the mechanisms of new word production. First, the amount of new words reflects how much the participants memorise things, particularly long-term memory. Theoretically, it was reported that Heaps’ law is related to the size of potential words^37^. Second, the number of new words suggests an ability of cognitive functions to take new information into their memory and to use it, which could be crucial especially for elderly people. People must cope with complex and unpredictable environments, but this ability could decline with aging.

Some of the scaling laws observed in brain and behaviour has been reported to be associated with functions or adaptability. For example, the movement patterns of various animals, including humans, often follow a scaling law, which can result in an efficient searching strategy for unpredictable environments^21^. For another example, it has been observed that the brain dynamics poised at a critical point between order and disorder follows a scaling law^38^, which makes it possible to achieve decision-making with flexibility and stability. Moreover, remarkably, theoretical studies on scaling laws of language show that Zipf’s law in language could be a consequence of the optimisation process for communication and feature of open-endedness, which refers to unbounded increasing complexity^29,33,39^. Our result shows that the deviations of exponents, that is, lower Heaps’ exponent, might imply a deviation from desirable and healthy states in cognitive functions. Therefore, it is crucial to understand the mechanisms of brain aging based on the difference between a healthy state and an impaired state in cognition in terms of scaling laws.

An intervention to improve cognitive functions is urgently needed in our aging society where disease-related aging is a serious problem. However, the effective strategies have not been understood fully. To come up with effective strategies, it is necessary to understand the mechanisms of cognitive functions and behaviour such as speaking and the causal relationship between them. In the present study, we revealed the association between cognitive functions and word production patterns and confirmed this using computational modelling. Bringing new information into spoken words may help to maintain and improve cognitive functions^40^. Next, it is important to understand how cognitive functions are improved through conversation or acquiring new words and using them.

## Methods

### Data collection

To obtain the data of spoken language from healthy elderly people, we recorded conversations among the participants. The participants were recruited from the Tokyo Silver Human Resources Center and were community-living healthy Japanese retired adults who speak Japanese as their mother-language. To limit participants to healthy people, the exclusion criteria were set as follows: dementia; neurological impairment; any disease or medication known to affect the central nervous system; and the Japanese Mini-Mental State Examination (MMSE-J) score of less than 24 points, which are the often-used criteria for screening dementia^41^. The assessment and screening to check participant eligibility were conducted based on medical interviews, neuropsychological tests, and self-reported questionnaires. Seventy-two people received the screening and seven were removed. Thus, the sample size for our data was 65 (30 males and 35 females). The mean age and range were 72.6 ± 3.2 (SD) and [66, 81], respectively. The MMSE-J score was 28.0±1.46 (mean±SD).

Before recording the conversations, we conducted cognitive functions tests for each participant. The tests included the Japanese version of Montreal Cognitive Assessment (MoCA-J) scores^42^, logical memory test (I + II) in WMS-R^43^, and the digit symbol coding test and digit span (forward + backward) in Wechsler Adult Intelligence Scale Third Edition (WAIS-III)^44^. MoCA-J was used to evaluate global cognitive function. Logical memory test I assesses immediate recall of the content of a story immediately after the examiner reads it, while logical memory test II assesses delayed recall 30 minutes later. The digit symbol coding test assesses the process speed and memory in digit symbol coding performance, which requires the subject to write down each corresponding symbol as fast as possible. The digit span (forward) assesses simple memory span, and digit span (backward) assesses working memory capacity. The RIKEN Institutional Review Board approved this study. All participants provided written informed consent. The characteristics of these values are shown in Supplementary Table 1.

Our recording of the conversations took place from June to September 2018 in an experimental room. Before the first recording, we divided the 65 participants into 16 groups. Fifteen groups had four participants and one group had five participants, based on participants’ availability. Every week, the participants joined the conversations for approximately 30 minutes, and the experiment sessions lasted for 14 weeks. Therefore, we obtained approximately seven hours’ conversational data from each group. The group members were fixed in their initially allocated groups until the end of all the experiments. In other words, each participant had conversations with the same group members each week. The 16 groups were divided into free conversation conditions and discussion conditions. The participants in eight of the 16 groups talked with each other freely. The participants in the other eight groups made a short presentation on a pre-determined theme (e.g. favourite places in the neighbourhood), which was given in advance each week and included Q&A sessions for participants within the same group. The latter is a method that we had developed previously for preventing dementia^40^, but here we focused not on the detail and effect of the method, but on the conversational patterns extracted from recorded conversational data.

### Conversational data pre-processing

To investigate word production patterns, we quantitatively analysed conversation transcriptions derived from the recorded audio data. For the analysis, we first applied Google Cloud Speech-to-Text (Google, Mountain View, CA, 2018) to automatic transcription from audio to text data, and then manually checked all the text by comparing it to the audio data and fixing any mistakes. Second, we automatically decomposed all text into words using MeCab (ver. 0.996), which is a useful tool for Japanese morphological analysis based on conditional random fields^45^. Finally, we obtained the data of the words that each participant spoke and used all the data in our analysis by putting together all the sessions, from the first to the fourteenth sessions. These analyses were conducted using R (ver. 3.4.3).

### Cognitive function scores

We conducted a principal component analysis to summarise four cognitive function scores of MoCA-J, WAIS III logical memory I + II (delayed), digit symbol coding, and digit span (forward + backward). The first principal component (PC1) contains 40.6% of all variances, and the coefficients of each cognitive score on PC1 have the same sign because the four cognitive function scores positively correlated with each other. Therefore, we can conclude that the larger the value on the PC1, the better the cognitive function. Hereafter, we use the value on the PC1 as ‘cognitive function score’. Additionally, for the simplicity of the interpretation, the cognitive score was normalised with mean = 0 and SD = 1.

### Zipf’s law and Heaps’ law

To quantitatively analyse word production patterns, we pay attention to scaling laws in language, which have been investigated in the context of statistical linguistics^16^. Previous studies have reported that most language data including corpus data robustly follow Zipf’s law and Heaps’ law^25,27,28,46^. Zipf’s law states that the relationship between the rank *r* of number of words and the frequency *P*(*r*) of words is described as *P*(*r*) ∼ *r*^−*α*^ where *α* is the scaling exponent and has been reported to be approximately 1, and “∼” means that the left side is proportional to the right side. Heaps’ law is about how the number of new words grows and that the relationship between the number of words *M* and the number of different words *N* follows a function *N* ∼ *M*^*β*^. In this study, we focused on whether words in the spoken language of healthy elderly people follow scaling laws and on the variation of exponents in the scaling relationships if scaling relationships exist.

For fitting the distribution to the data of rank-frequency relationship, we compared seven candidate distributions including a power-law, shifted power-law, cutoff power-law, and so on (see the details under Supplementary Note). First, we fitted each candidate model to the data using maximum likelihood estimation (MLE)^34^ and estimated the parameters (i.e. exponents) using the Nelder-Mead method for maximising the log-likelihood. Then, we explored the best model using AICs and Akaike weights. As for Heaps’ law, we used the least-squared method, estimated the scaling exponents and calculated the R-squared value to evaluate good-of-fit.

### A mathematical model for scaling law and cognitive functions

To investigate how the variations of the scaling exponents in Heaps’ law emerge from the difference of cognitive functions or behavioural rules, we used a mathematical generative model based on a previously proposed stochastic model for book corpus data^25^. In the model, new words can be produced by the following rules. When a word is produced, a novel word is produced at the probability *p*_new_ and the already used word is used at the probability 1 − *p*_new_. The probability *p*_new_ is updated depending on a function of *n*_c_, which is the number of the different words every time a novel word is produced. The update rule is as follows:

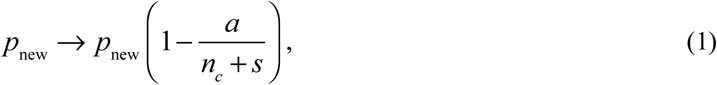

where *a* is decay parameter and *s* is a small constant, and can produce the scaling law of Heaps’ law. Here, we interpret *a* as 1/*γ*, where *γ* is a parameter of cognitive functions because *a* is directly leading to the exponent *β*^25^. Moreover, we observed the double scaling law in the relationship between the number of words and different words. In the case of *M* < *m*, *N* = *M*^1^ and for *M* ≥ *m*, *N* ∼ *M*^*β*^. To model this property, we adopted a different rule for small *M* following Gerlach and Altmann^25^. If *N* is small, here *N* < 32, the empirical mean value of break points *m*, we assumed that new words were produced at probability 0.8, but the higher value produced robust results. For *N* ≥ 32, we used the equation (1). The two rules made it possible to switch from a linear relationship to a sublinear relationship.

## Acknowledgement

We thank Dr. Takuya Sekiguchi and Dr. Yukie Sano for fruitful discussions. We also thank Dr. Seiki Tokunaga, Megumi Kubota, and Kaai Yamaguchi for helping with data collection. This study was supported by JSPS KAKENHI Grant Number JP18K18140.

## References

1. Wimo, A. et al. The worldwide costs of dementia 2015 and comparisons with 2010. Alzheimers Dement. 13, 1–7 (2017).

2. Livingston, G. et al. Dementia prevention, intervention, and care. The Lancet 390, 2673–2734 (2017).

3. Huntley, J. D. & Howard, R. J. Working memory in early Alzheimer’s disease: a neuropsychological review. Int. J. Geriatr. Psychiatry 25, 121–132 (2010).

4. Cerejeira, J., Lagarto, L. & Mukaetova-Ladinska, E. B. Behavioral and Psychological Symptoms of Dementia. Front. Neurol. 3, (2012).

5. Taler, V. & Phillips, N. A. Language performance in Alzheimer’s disease and mild cognitive impairment: A comparative review. J. Clin. Exp. Neuropsychol. 30, 501–556 (2008).

6. Szatloczki, G., Hoffmann, I., Vincze, V., Kalman, J. & Pakaski, M. Speaking in Alzheimer’s Disease, is That an Early Sign? Importance of Changes in Language Abilities in Alzheimer’s Disease. Front. Aging Neurosci. 7, (2015).

7. Fraser, K. C., Meltzer, J. A. & Rudzicz, F. Linguistic Features Identify Alzheimer’s Disease in Narrative Speech. J. Alzheimers Dis. 49, 407–422 (2015).

8. Hernández-Domínguez, L., Ratté, S., Sierra-Martínez, G. & Roche-Bergua, A. Computer-based evaluation of Alzheimer’s disease and mild cognitive impairment patients during a picture description task. Alzheimers Dement. Diagn. Assess. Dis. Monit. 10, 260–268 (2018).

9. Asgari, M., Kaye, J. & Dodge, H. Predicting mild cognitive impairment from spontaneous spoken utterances. Alzheimers Dement. Transl. Res. Clin. Interv. 3, 219–228 (2017).

10. Hauser, M. D., Chomsky, N. & Fitch, W. T. The Faculty of Language: What Is It, Who Has It, and How Did It Evolve? Science 298, 12 (2002).

11. Michel, J.-B. et al. Quantitative Analysis of Culture Using Millions of Digitized Books. Science 331, 176–182 (2011).

12. Monsch, A. U. et al. Comparisons of Verbal Fluency Tasks in the Detection of Dementia of the Alzheimer Type. Arch. Neurol. 49, 1253–1258 (1992).

13. Snowdon, D. A. Linguistic ability in early life and cognitive function and Alzheimer’s disease in late life. Findings from the Nun Study. JAMA J. Am. Med. Assoc. 275, 528–532 (1996).

14. Fraser, K. C., Lundholm Fors, K. & Kokkinakis, D. Multilingual word embeddings for the assessment of narrative speech in mild cognitive impairment. Comput. Speech Lang. 53, 121–139 (2019).

15. Bak, P. How nature works: the science of self-organized criticality. (Copernicus, 1996).

16. Zipf, G. K. Human behavior and the principle of least effort. (Addison-Wesley, 1949).

17. Barabási, A.-L. The origin of bursts and heavy tails in human dynamics. Nature 435, 207–211 (2005).

18. Cattuto, C., Loreto, V. & Pietronero, L. Semiotic dynamics and collaborative tagging. Proc. Natl. Acad. Sci. 104, 1461–1464 (2007).

19. Kello, C. T. et al. Scaling laws in cognitive sciences. Trends Cogn. Sci. 14, 223–232 (2010).

20. Mora, T. & Bialek, W. Are Biological Systems Poised at Criticality. J. Stat. Phys. 144, 268–302 (2011).

21. Viswanathan, G. M., Da Luz, M. G., Raposo, E. P. & Stanley, H. E. The physics of foraging: an introduction to random searches and biological encounters. (Cambridge University Press, 2011).

22. Proekt, A., Banavar, J. R., Maritan, A. & Pfaff, D. W. Scale invariance in the dynamics of spontaneous behavior. Proc. Natl. Acad. Sci. 109, 10564–10569 (2012).

23. Karsai, M., Jo, H. H. & Kaski, K. Bursty human dynamics. (Springer International Publishing, 2018).

24. Jin, C., Song, C., Bjelland, J., Canright, G. & Wang, D. Emergence of scaling in complex substitutive systems. Nat. Hum. Behav. 3, 837–846 (2019).

25. Gerlach, M. & Altmann, E. G. Stochastic Model for the Vocabulary Growth in Natural Languages. Phys. Rev. X 3, 021006 (2013).

26. Bian, C., Lin, R., Zhang, X., Ma, Q. D. Y. & Ivanov, P. Ch. Scaling laws and model of words organization in spoken and written language. EPL Europhys. Lett. 113, 18002 (2016).

27. Piantadosi, S. T. Zipf’s word frequency law in natural language: A critical review and future directions. Psychon. Bull. Rev. 21, 1112–1130 (2014).

28. Heaps, H. S. Information retrieval, computational and theoretical aspects. (Academic Press, 1978).

29. Cancho, R. F. i. & Sole, R. V. Least effort and the origins of scaling in human language. Proc. Natl. Acad. Sci. 100, 788–791 (2003).

30. Font-Clos, F. & Corral, Á. Log-Log Convexity of Type-Token Growth in Zipf’s Systems. Phys. Rev. Lett. 114, 238701 (2015).

31. Piotrowskii, R. H. Statistical models of text and their linguistic and synergetic analysis. Autom. Doc. Math. Linguist. 41, 159–170 (2007).

32. Baixeries, J., Elvevåg, B. & Ferrer-i-Cancho, R. The Evolution of the Exponent of Zipf’s Law in Language Ontogeny. PLoS ONE 8, e53227 (2013).

33. Ferrer i Cancho, R. The variation of Zipf?s law in human language. Eur. Phys. J. B 44, 249–257 (2005).

34. Clauset, A., Shalizi, C. R. & Newman, M. E. J. Power-Law Distributions in Empirical Data. SIAM Rev. 51, 661–703 (2009).

35. Yule, G. U. A Mathematical Theory of Evolution, Based on the Conclusions of Dr. J. C. Willis, F.R.S. Philos. Trans. R. Soc. Lond. Ser. B Contain. Pap. Biol. Character 21, 21–87 (1925).

36. Calude, A. S. & Pagel, M. How do we use language? Shared patterns in the frequency of word use across 17 world languages. Philos. Trans. R. Soc. B Biol. Sci. 366, 1101–1107 (2011).

37. Sano, Y., Takayasu, H. & Takayasu, M. Zipf’s Law and Heaps’ Law Can Predict the Size of Potential Words. Prog. Theor. Phys. Suppl. 194, 202–209 (2012).

38. Chialvo, D. R. Emergent complex neural dynamics. Nat. Phys. 6, 744–750 (2010).

39. Corominas-Murtra, B., Seoane, L. F. & Solé, R. Zipf’s Law, unbounded complexity and openended evolution. J. R. Soc. Interface 15, 20180395 (2018).

40. Otake-Matsuura, M. et al. Photo-Integrated Conversation Moderated by Robots for Cognitive Health in Older Adults: A Randomized Controlled Trial. Preprint at medRxiv https://www.medrxiv.org/content/10.1101/19004796v1 (2019).

41. Sugishita, M. et al. The validity and reliability of the japanese version of the mini-mental state examination (MMSE-J) with the original procedure of the attention and calculation task. Japanes J. Cogn. Neurosci. 20(2), 91–110.

42. Fujiwara, Y. et al. Brief screening tool for mild cognitive impairment in older Japanese: Validation of the Japanese version of the Montreal Cognitive Assessment: Brief screening tool for MCI. Geriatr. Gerontol. Int. 10, 225–232 (2010).

43. Wechsler, D. A. Wechsler Memory Scale-Revised. San Antonio: The Psychological Corporation (1987).

44. Wechsler, D. A. Wechsler Adult Intelligence Scale - Third edition. San Antonio: The Psychological Corporation (1997).

45. Kudo, T., Yamamoto, K. & Matsumoto, Y. Applying conditional random fields to Japanese morphological analysis. Proceedings of the 2004 Conference on Empirical Methods in Natural Language Processing, 230–237 (2004).

46. Smith, R. Investigation of the Zipf-Plot of the Extinct Meroitic Language. Preprint at arXiv https://arxiv.org/abs/0808.2904 (2008).

