## Supplemental Note for "Scaling laws in spoken language associated with cognitive functions"

### Supplementary note for
### “Scaling laws in spoken language associated with cognitive functions”

Masato S. Abe[1] and Mihoko Otake-Matsuura[1]

1. RIKEN Center for Advanced Intelligence Project, Nihonbashi 1-chome Mitsui Building, 15th floor, 1-4-1 Nihonbashi, Chuo-ku, Tokyo 103-0027, Japan (Tel: +81-048-467-3627)

#### 1. Basic characteristics for words

In our study, we conducted recordings of conversations among healthy elderly people, and the length of total recordings was approximately seven hours for each group. The conversations did not break off in all the data. Therefore, we obtained recording data of about 1.75 hours for each participant on average because each group had four participants, except for one group that had five participants. However, there were variations among participants in the number of words spoken. The distributions of words and different words are shown in Supplementary Fig. 1. Some participants were talkative, while others were not. Our main result for the association between cognitive scores and the scaling exponent (Fig. 2a) was obtained from all data sets.

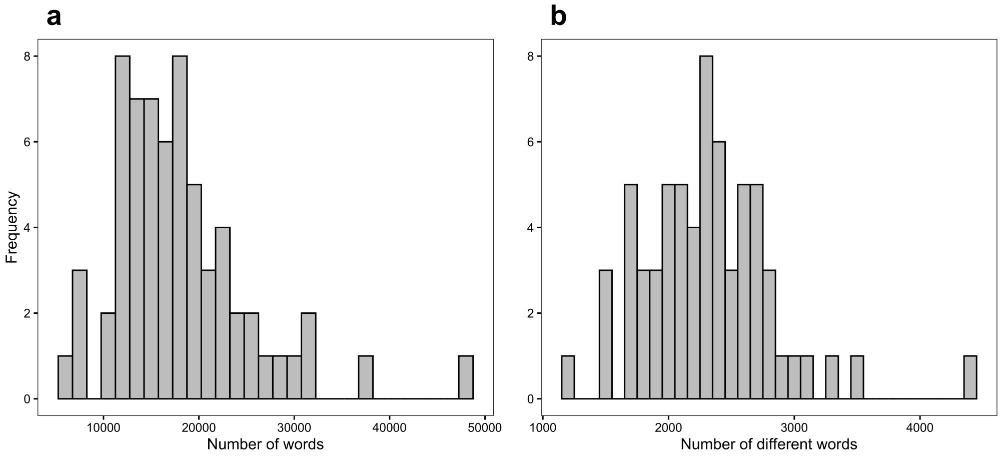


Supplementary Figure 1: The distributions of the number of words and different words

Supplementary Table 1: Cognitive scores

| | MoCA-J | Logical memory (I + II) | Digit symbol coding | Digit span (forward + backward) |
|---|---|---|---|---|
| Mean ± SD | 25.7 ± 2.6 | 16.7 ± 6.9 | 53.8 ± 13.4 | 16.0 ± 3.0 |

#### 2. Fitting procedure for Zipf's distribution

We used rigorous statistical methods for fitting a probability distribution to empirical rank-word frequency relationships[1,2]. Here, the candidate distributions included a power-law distribution, a shifted power-law distribution, a power-law with exponential cutoff (tail), a log-normal distribution, Weibul distribution, and a double power-law distribution. The probability mass functions for such distributions are shown in Supplementary Table 1. We estimated the parameters of each distribution numerically by maximising log-likelihood. This procedure was conducted using the Nelder-Mead method implemented in the optim function of R. The constant $C$ is the value for normalising and we obtained it as $\sum_{r=1}^{N_{\max}} F(r;\, U) = 1$, where $U$ is a set of parameters, and $N_{\max}$ is the maximal number of different words for each participant.

Supplementary Table 2: Probability mass functions for each fitting model

| Distribution | Probability mass function $F(r; U)$ |
|---|---|
| Power-law | $Cr^{-\alpha}$ |
| Shifted power-law | $C(r+b)^{-\alpha}$ |
| Power-law with exponential cutoff (tail) | $C\exp(-br)r^{-\alpha}$ |
| Log-normal | $Cr^{-1}\exp(-0.5(\ln r - \mu)^2 / \sigma^2)$ |
| Weibull | $Cr^{\alpha-1}\exp(-br^{-\alpha})$ |
| Double power-law | $C\begin{cases} r^{-1}, & (r \le b) \\ b^{\alpha-1} r^{-\alpha} & (r > b) \end{cases}$ |

After estimating the parameters using MLE, we calculated Akaike Information Criteria (AIC), which is defined as

$$\mathrm{AIC}_i = -2(\text{log-likelihood of model } i) + 2(\text{number of parameters in model } i).$$

Then, to compare these distributions in terms of model selection, the Akaike weights= $w_i$ of fitting model $i$ was calculated following[3]:

$$w_i = \frac{\exp\left[-(\mathrm{AIC}_i - \mathrm{AIC}_{\min})/2\right]}{\sum_{j=1}^{R}\left\{\exp\left[-(\mathrm{AIC}_j - \mathrm{AIC}_{\min})/2\right]\right\}},$$

where $R$ is the number of candidate models, here 6, and $\mathrm{AIC}_{\min}$ is the minimum AIC in those of fitting models. The value ranges from 0 to 1 and represents the probability of the model given the data[3]. Therefore, the model producing $w_i$ close to 1 gives us the evidence of supporting the model. Supplementary Fig. 2 illustrates an example of a fitting result of a particular participant.

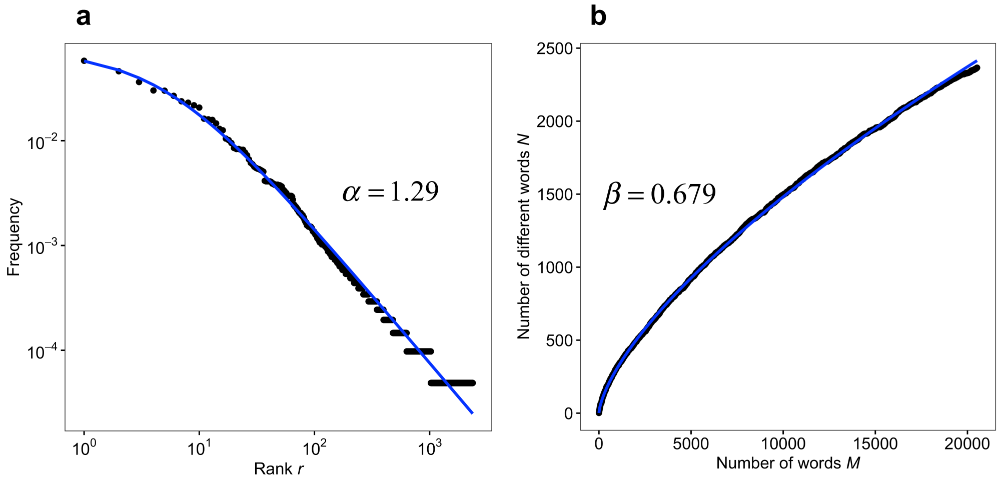


Supplementary Figure 2: Examples of fitted distribution and function to data
The black dots represent empirical data for a participant, and the blue lines are fitted distribution for the rank-frequency relationship (a) and for the relationship between the number of words and distinct words (b).

#### 3. Relationship between scaling exponents and raw cognitive scores

In the main text, we reported the relationship between the scaling exponents and the first component derived from the principal component analysis of four cognitive scores. Here, we show the relationship between the scaling exponents and the four raw cognitive scores. Supplementary Table 2 suggests that all the four cognitive scores are associated with the scaling exponents $\beta$ of Heaps' law.

Supplementary Table 3: Regression analysis on scaling exponents

The results are derived from an analysis of all data sets.

| | Explanatory variable: Estimates (SE, $p$-value) | | | | |
|---|---|---|---|---|---|
| | Cognitive score | Gender (male) | Condition (presentation) | Age | Constant |
| Scaling exponent $\alpha$ | 0.011 (0.006, 0.079) | 0.020 (0.012, 0.084) | 0.011 (0.012, 0.351) | 0.002 (0.002, 0.334) | −1.362** (0.134, $1.1 \times 10^{-14}$) |
| Scaling exponent $\beta$ | 0.008** (0.003, 0.002) | 0.011* (0.005, 0.033) | 0.002 (0.005, 0.750) | 0.0003 (0.001, 0.722) | 0.658** (0.058, $< 2 \times 10^{-16}$) |

*$p < 0.05$; **$p < 0.01$

Supplementary Table 4: Correlation coefficients between each raw cognitive score and the scaling exponent $\beta$

The results are derived from an analysis of all data sets.

| Cognitive score | Correlation coefficient $r$ | $p$-value |
|---|---|---|
| MoCA-J | 0.37** | 0.002 |
| Logical memory (I + II) | 0.30* | 0.015 |
| Digit symbol coding | 0.32** | 0.009 |
| Digit span (forward + backward) | −0.13 | 0.315 |

*$p < 0.05$; **$p < 0.01$

### References

1. Gerlach, M. & Altmann, E. G. Stochastic Model for the Vocabulary Growth in Natural Languages. *Phys. Rev. X* **3**, 021006 (2013).
2. Clauset, A., Shalizi, C. R. & Newman, M. E. J. Power-Law Distributions in Empirical Data. *SIAM Rev.* **51**, 661–703 (2009).
3. Burnham, K. P., & Anderson, D. R. Multimodel inference: understanding AIC and BIC in model selection. *Sociol. Methods Res.* **33**(2), 261-304 (2004).
